# Open-Source 3*D* Active Sample Stabilization for Fluorescence Microscopy

**DOI:** 10.1101/2025.01.10.631718

**Authors:** Sanket Patil, Giuseppe Vicidomini, Eli Slenders

## Abstract

Super-resolution microscopy has enabled imaging at nanometer-scale resolution. However, achieving this level of detail without introducing artifacts that could mislead data interpretation requires maintaining sample stability throughout the entire imaging acquisition. This process can range from a few seconds to several hours, particularly when combining live-cell imaging with super-resolution techniques. Here, we present a 3*D* active sample stabilization system based on real-time tracking of fiducial markers. To ensure broad accessibility, the system is designed using readily available off-the-shelf optical and photonic components. Additionally, the accompanying software is open-source and written in Python, facilitating adoption and customization by the community. We achieve a standard deviation of the sample movement within 1 nm in both the lateral and axial direction for a duration in the range of hours. Our approach allows easy integration into existing microscopes, not only making prolonged super-resolution microscopy more accessible but also allowing confocal and widefield live-cell imaging experiments spanning hours or even days.

## Introduction

Over the past three decades, super-resolution microscopy (SRM) techniques have emerged, enabling a spatial resolution far beyond the diffraction limit (1–5). Notably, in the context of single-molecule localization microscopy, resolutions at the Ångström scale have recently been achieved (6–8). SRM opened up the possibility to study cellular processes at the nm scale, allowing *e*.*g*., nanoscale functional topography of chromatin and studying its role in genome integrity (9, 10), and the direct visualization of kinesin dynamics (11, 12). However, the performance of SRM, particularly of Single-Molecule Localization Microscopy (SMLM), depends on the inherent stability of the whole imaging system, including the sample. System drift, a pervasive challenge affecting all imaging platforms, significantly hinders the achievable resolution in SRM. This drift arises from a combination of thermal fluctuations and mechanical vibrations within both the imaging system itself and the surrounding environment. The negative impact of system drift becomes evident in the form of a declining image quality ranging from minor imperfections in low-resolution imaging to severe spatial resolution degradation, potential misinterpretation of the data, and even rendering the data unusable in super-resolution applications.

Approaches to address sample drift can be broadly categorized into two methods: active stabilization (13–15) and post-processing corrections (16–18). Active stabilization focuses on real-time drift correction by continuously monitoring the sample’s position, for example, by tracking fiducial markers, and by using mechanical or piezoelectric actuators to adjust the sample’s position to counteract any drift. This approach can be implemented for all three axes or for axial direction only in scenarios where lateral drift is negligible or addressed through separate corrections (19). Post-processing corrections address drift after image acquisition. Drift estimation can be achieved either through mathematical model estimation (20) or by imaging fiducial markers alongside the sample (13, 15). Notably, without acquiring a full 3D image stack, axial drift cannot be effectively compensated during post-processing. Indeed, axial drift leads to out-of-focus images, with inherent information loss as a consequence. Active stabilization is preferred in low-throughput SMLM techniques, such as MINFLUX, RAST-MIN, and pMINFLUX (21–24), where the limited number of localized molecules makes it difficult to accurately deduce drift through post-processing methods.

The use of active stabilization techniques to enhance sample stability has become more widespread in recent years (19, 25–28). Additionally, multiple commercial systems with axial stabilization capabilities have emerged (29– 31). However, the adaptability of these commercial solutions is often hindered by their limited compatibility with custom-built imaging systems. This has led to the development of numerous custom approaches for sample stabilization, broadly categorized into fiducial marker-based stabilization and reflection-based stabilization. Fiducial marker-based stabilization tracks the position of pre-introduced fiducial markers, such as gold nanoparticles, gold nanorods, or fluorescent beads, within the sample. Drift is continuously monitored through a dedicated imaging pathway that tracks in real-time the position of these markers. Reflection-based stabilization makes use of the reflection of a focused beam off the coverslip. An axial movement of the sample leads to a change in the position or shape of the image of the reflected beam. In both cases, a feedback loop adjusts the sample position to actively compensate for any detected displacement. Each approach presents distinct advantages and limitations. Fiducial-based techniques offer the benefit of three-dimensional stabilization but necessitate the introduction of fiducial markers during sample preparation. Conversely, reflection-based techniques, while requiring no sample alteration, are restricted only to axial correction. Consequently, fiducial-based stabilization strategies are often the preferred choice for low throughput SMLM methods, such as MINFLUX, which require real-time 3*D* stabilization.

Several active 3*D* stabilization systems for custom-built microscopes currently exist (22, 24, 32). While these approaches yield stabilization results around 1 nm for all axes, their implementation is either hindered by the need for a costly setup, including a PC with a suitable graphics card for GPU computing, a more complex sample preparation, such as the use of functionalized polystyrene beads, a limited feedback update rate (< 20 Hz), or the lack of opensource availability. Here, we present a fast, open-source 3D active sample stabilization system that delivers similar performance but at a low cost, requiring neither complex sample preparation nor extensive hardware installation. The setup is designed as a modular, standalone add-on that is compact and can be easily integrated with any microscope. Besides the actuators to move the sample, the hardware is based on readily available commercial components, with a total implementation cost around C2500.

## Materials and methods

### A Optical Setup and sample

The optical configuration is shown in Fig. 1a. An infrared laser (L780P010 - 780 nm, 10 mW, Thorlabs, New Jersey, USA) serves as the illumination source for the fiducial markers. The collimated beam is split by a 50/50 cube beam splitter (BS014, 700 - 1100 nm, Thorlabs) and coupled into the microscope with a near-infrared (NIR) short-pass dichroic mirror (AHF 745/SP BrightLine HC Shortpass Filter, F39-745). By adjusting the mirrors positioned before the beam splitter, the beam is laterally displaced from the optical axis at the back aperture of the objective, leading to highly inclined and laminated optical sheet (HiLo) illumination after passing through the tube lens and objective lens (Nikon Plan Apo VC 100*×*/1.40 Oil OFN25 DIC N2). The HiLo illumination enables imaging over a large FOV (∼36 *×* 36 µm), which allows tracking of multiple particles simultaneously to improve robustness and precision (in our experiments, we use a concentration providing a number of particles between 5 and 15). The backscattered signal from the fiducial markers is collected by the same objective and reflected by the NIR dichroic mirror to enter the stabilization part again. Here, the beam passes through the 50/50 cube beamsplitter and enters a 4f system composed of two lenses, *L*_1_ and *L*_2_ (f = 200 mm each). Both the tube lens and *L*_1_, and *L*_1_ and *L*_2_ form a 4f system. In the conjugate plane between *L*_1_ and *L*_2_, an image of the back focal plane is formed, and the light reflected by the cover slip is focused off-axis. An iris diaphragm (IDA25/M, Thorlabs) blocks the light reflected by the coverslip while transmitting most of the backscattered light. Finally, the signal is split with another 50/50 beam splitter and projected onto two CMOS cameras (CS165MU/M - Zelux, Thorlabs), used to independently stabilize the lateral and ax-ial direction, respectively. A cylindrical lens (f = 1000 mm) induces astigmatism into the latter beam path to enable axial position estimation. Using two cameras instead of one effectively prevents aberrations caused by the cylindrical lens from introducing errors in lateral position calculations. The captured images from both cameras are transmitted to a central computer for further processing and calculations. Drift corrections are sent to a three-axis nanopositioning piezoelectric stage (P-545.3R8S PInano, E-727 Controller, Physik Instrumente).

**Fig 1.**
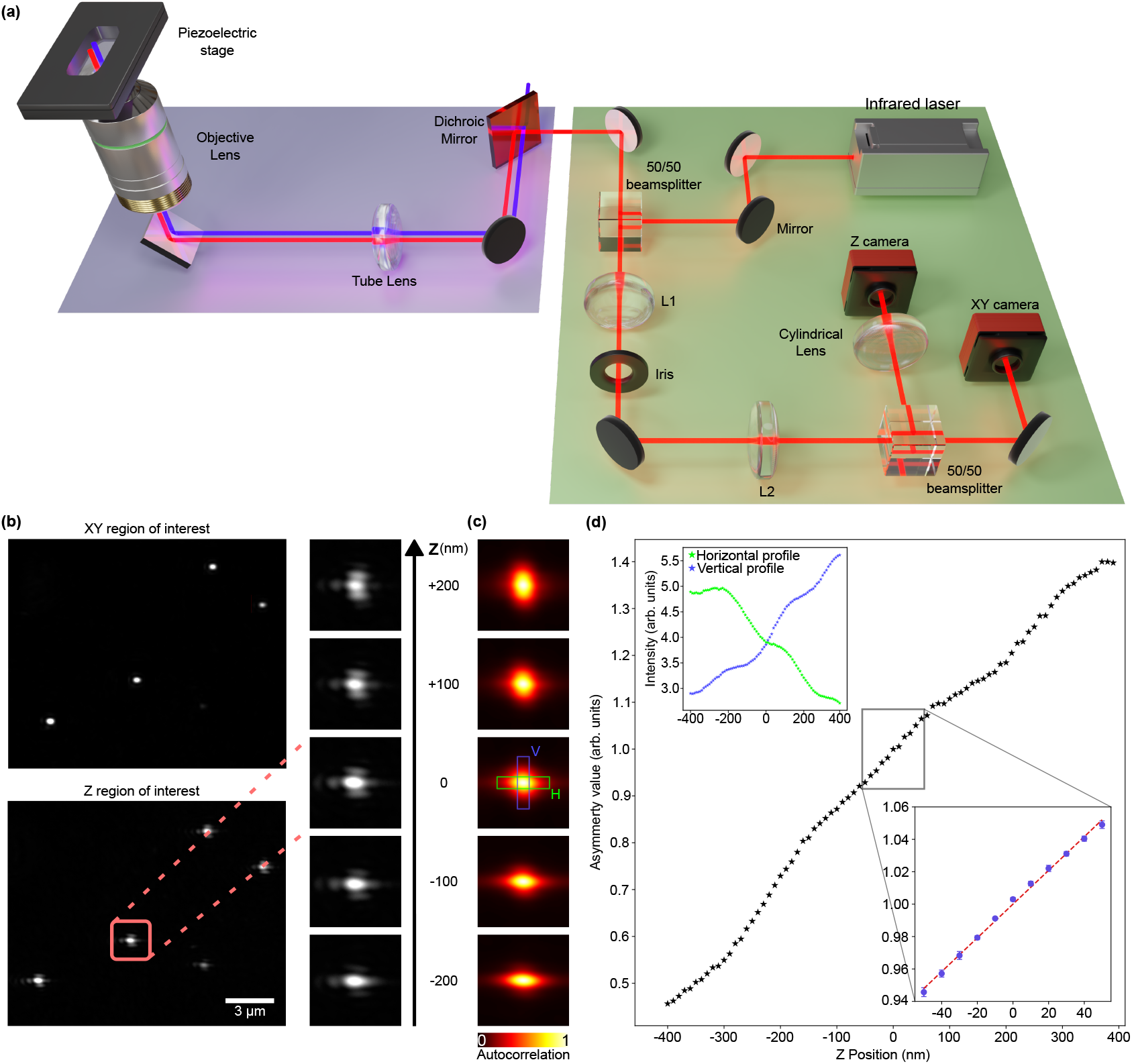
Optical Setup and Axial Calibration. (a) Optical setup of the stabilization system. The green part shows the stabilization setup, which is coupled to a microscope (partially shown in purple). The stabilization beam is coupled into the imaging path using a short-pass dichroic mirror, depicted in red in the imaging part. (b) Exemplary camera images of the XY and Z camera. The column on the right shows the astigmatism induced by the cylindrical lens, as imaged by the Z camera at different axial planes. (c) Autocorrelation of the images captured by the Z camera for the same z positions as in (b). The axial position of the sample is derived from the asymmetry of the autocorrelation function, calculated as the ratio of the summed intensities of the pixel values in H (green) and V (blue). (d) Measured autocorrelation asymmetry as a function of the *z* position. The calibration measurement performed at the start of each experiment covers a smaller range (from −50 nm to +50 nm, see inset), where the asymmetry can be well approximated as a linear curve (dotted line). The calibration curve and the error bars are the mean and standard deviation over ten iterations. The inset plot in the top left shows the horizontal and vertical components (shown in (c)) of the summed intensities over the green and blue boxes.

To stabilize the sample, fiducial markers such as gold nanoparticles (GNPs) can be used. Here, we use GNPs with a diameter ranging from 100-150 nm. The choice of gold nanoparticles stems from their several advantageous characteristics: readily available and cost-effective at various sizes, and capable of scattering laser light across a wide range of wavelengths (33). The components required for building the stabilization are listed in Table S1 and Table S2.

### B Control Software

The system integrates a central computer interfacing with the CMOS cameras and the nanopositioning piezoelectric stage. The computer stabilizes the sample by imaging the fiducial markers, performing real-time 3*D* locaisation of the particles, and repositioning the stage accordingly. The software is developed in Python and runs entirely on the CPU. We use multiprocessing to parallelize fiducial marker localization in the lateral direction, accelerating drift calculations. The decision to forgo GPU utilization ensures broad compatibility across a diverse range of systems, including systems equipped solely with integrated GPUs or non-NVIDIA dedicated options, where GPU utilization might not offer a performance advantage and could even impede processing speed due to data transfer overhead.

Before initiating the stabilization process, the software connects to the two CMOS cameras and the nanopositioning piezoelectric stage. All three pieces of hardware require dynamic-link library (.dll) files to establish a connection using the USB 3.0 protocol. These .dll files, provided by the proprietary control software of the cameras (Thorlabs) and the piezoelectric stage (Physik Instrumente), can be installed by the user via a straightforward Windows installation wizard.

After the hardware connection, the user can crop the FOV for each camera independently, Fig. 1 (b), balancing a larger FOV to capture more particles with a smaller FOV to achieve a higher frame rate. Subsequently, the user selects a chosen number of particles within this FOV for lateral stabilization. The software then performs a *z*-scan using the same piezoelectric stage employed for stabilization to measure the PSF shape at various axial positions, Fig. 1 (b).

Then, depending on the selected settings, up to 10 processes start running in parallel: one process per camera for image capture, one process per camera for image analysis, one process to control the stage, one process for real-time plotting, and up to four additional processes for miscellaneous tasks, such as data saving, statistical calculations, and data logging. This implementation accommodates processes running at different speeds, allowing capturing images at a high frame rate (*e*.*g*., ∼120 Hz), even when stage updates occur at a lower frequency (∼20 Hz). Fig. 2 illustrates the pseudocode for the stabilization software. Of the 10 processes initiated, 5 are essential for actively stabilizing the sample, while the remaining 5 are optional and can be toggled on or off depending on experimental requirements. Disabling the non-essential processes can enhance acquisition and calculation speed when the CPU is fully utilized. When active, all 10 processes operate in parallel, utilizing a shared memory space allocated at the start of the program to facilitate simultaneous data access. In this way, the data generated by each process is shared with the other processes. Within this shared memory, data is classified into two types: new data (indicated by green arrows) generated with each process iteration and duplicate data (indicated by pink arrows), which is used for data saving and logging purposes. A potential challenge arises when multiple processes attempt to access the shared memory concurrently for reading or writing, which can lead to memory corruption and disrupt the stabilization program. To address this issue, we have implemented a memory lock mechanism. The memory lock is implemented in such a way that whenever a process needs to read from or write to the shared memory buffer, the memory is locked by that particular process handler so that no other process can access that memory space. While the memory lock introduces a minor time penalty of less than 100 µs per iteration, it enables efficient data exchange between processes without the need for explicit inter-process communication (IPC) mechanisms, ensuring stable and uninterrupted program operation.

**Fig 2.**
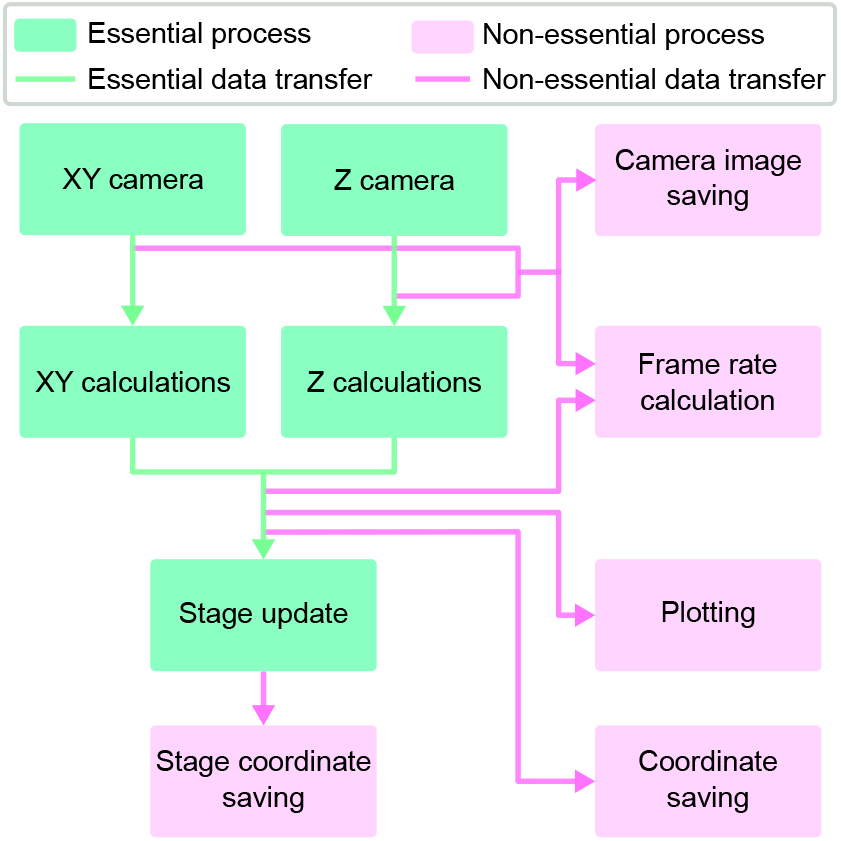
Stabilzaition code workflow. Flowchart with pseudocode describing the stabilization code that controls the hardware. The code is split into 10 different processes that are categorized into two types. The first type (green) represents the essential processes required for the stabilization code. The second type (pink) represents the non-essential part of the stabilization code. Similarly, the green arrows show the essential data transfer, and the pink arrows show the non-essential data transfer.

Positional deviations in 3*D* are assessed by comparing measured particle locations with the reference points established at the start of the measurement. Detected deviations trigger corresponding negative feedback commands sent to the piezoelectric stage. This closed-loop control system actively returns the sample to its relative position with respect to the stabilization beam path. In more detail, the software reads the cameras and stores the last 25 frames in memory. A separate process takes, for each camera, the weighted average of these frames, with the highest weight for the latest frame and an exponentially decreasing weight for earlier frames. For lateral drift correction, the particles are localized, and their average drift from the reference points is stored in memory. A separate process checks with a fixed rate (*×*20 per second) whether or not the measured average drift exceeds a threshold of 1 nm and, if necessary, updates the lateral position of the stage. Axial drift is assessed by comparing the PSF shape in the averaged frame with the reference curve obtained during the initialization procedure. Similarly to the lateral drift, a separate process updates the axial position of the stage with a fixed rate of 20 Hz. To prevent oscillatory behavior caused by overshooting, we mul-tiply the measured drift for each dimension with a factor *α* (0 *< α* ≤ 1). This gain reduction allows approaching the set point gradually, *i*.*e*., in multiple stage movement cycles, which improves the stability.

### C Position Estimation

We estimate the lateral drift in the camera frame of reference **d**(*t*) at time point *t* by averaging the measured changes in the 2*D* position of *N* fiducial markers compared to the start of the measurement: **d**(*t*) = ⟨x(*t*) − x(0)⟩, with x(*t*) the positions of the particles at time *t*. We use the local gradients method (34) to measure x(*t*) because of its computational efficiency compared to fitting a Gaussian function. Conversion of drift from pixels to nm is done based on a calibration measurement with a reflective grid sample (R1L3S3PR, ThorLabs).

Axial drift estimation is based on astigmatism introduced by the cylindrical lens positioned in front of the camera and comparing the shape of the point spread function (PSF) with respect to the start of the measurement. To estimate the relative *z*-position, we calculate the autocorrelation *G* in a user-selected region of the image, ideally containing multiple particles for enhanced robustness and precision (Fig. 1 (c, d)). We define a measure for the PSF shape *A* as the ratio *A* =Σ_*i*_ *G*_*i*_*/*Σ_*j*_ *G*_*j*_ with *i* ∈ *H* and *j* ∈ V, with *H* and *V* perpendicular rectangular regions centered around the autocorrelation peak. Conversion from *A* to axial displacement in nm is based on a reference curve measured at the start of every measurement (Fig. 1 (d)). This calibration involves sampling multiple planes in both the positive (+*z*) and negative (−*z*) directions with respect to the experiment plane. At each sampled plane, we calculate the corresponding *A* value. These values are then fitted using linear regression (see the inset of Fig. 1 (e)). The slope of the fit represents the conversion factor Δ*A/*Δ*z*, expressed in arbitrary units per nanometer. We typically perform the calibration procedure for a small range (−50 nm to +50 nm) with respect to the initial *z* position, where the calibration curve can be well approximated by a linear curve.

## Results and discussion

First, we measured the performance of the stabilization system using the calculated particle positions with respect to the start of the measurement in the camera frame of reference, as measured by the stabilization system itself during closed-loop stabilization (Fig. 3 (a)). For all axes, we found a normal distribution with a mean position around 0 and a standard deviation below 1 nm for a 1000 s measurement. Since the nanopositioning piezo system has a closed-loop resolution of 1 nm, and our stabilization software uses a 1 nm threshold for stage updates, we cannot expect a better precision. Note that, due to differences in the frame rate of the cameras and the update rate of the stage, the measured drift may be higher than the 1 nm threshold for multiple consecutive frames, as shown in the insets in Fig. 3 (a). The corresponding stage position (Fig. 3(b)), shows about 100 and 200 nm drift in the *x, y* direction, respectively, and about 150 nm drift in the axial direction over 1000 s. Comparison of the power spectral density curves of the stage and sample position (Fig.3(c)) shows that drift and low-frequency vibrations are suppressed by the stabilization software by more than one order of magnitude in all dimensions. The repositioning of the stage amplifies some vibrations, as can be seen around 1 Hz for the *x*-direction and 2.5 Hz for the *z*-direction. However, the amplitudes of these vibrations are low enough to keep the overall sample movement below 1 nm standard deviation in each direction.

**Fig 3.**
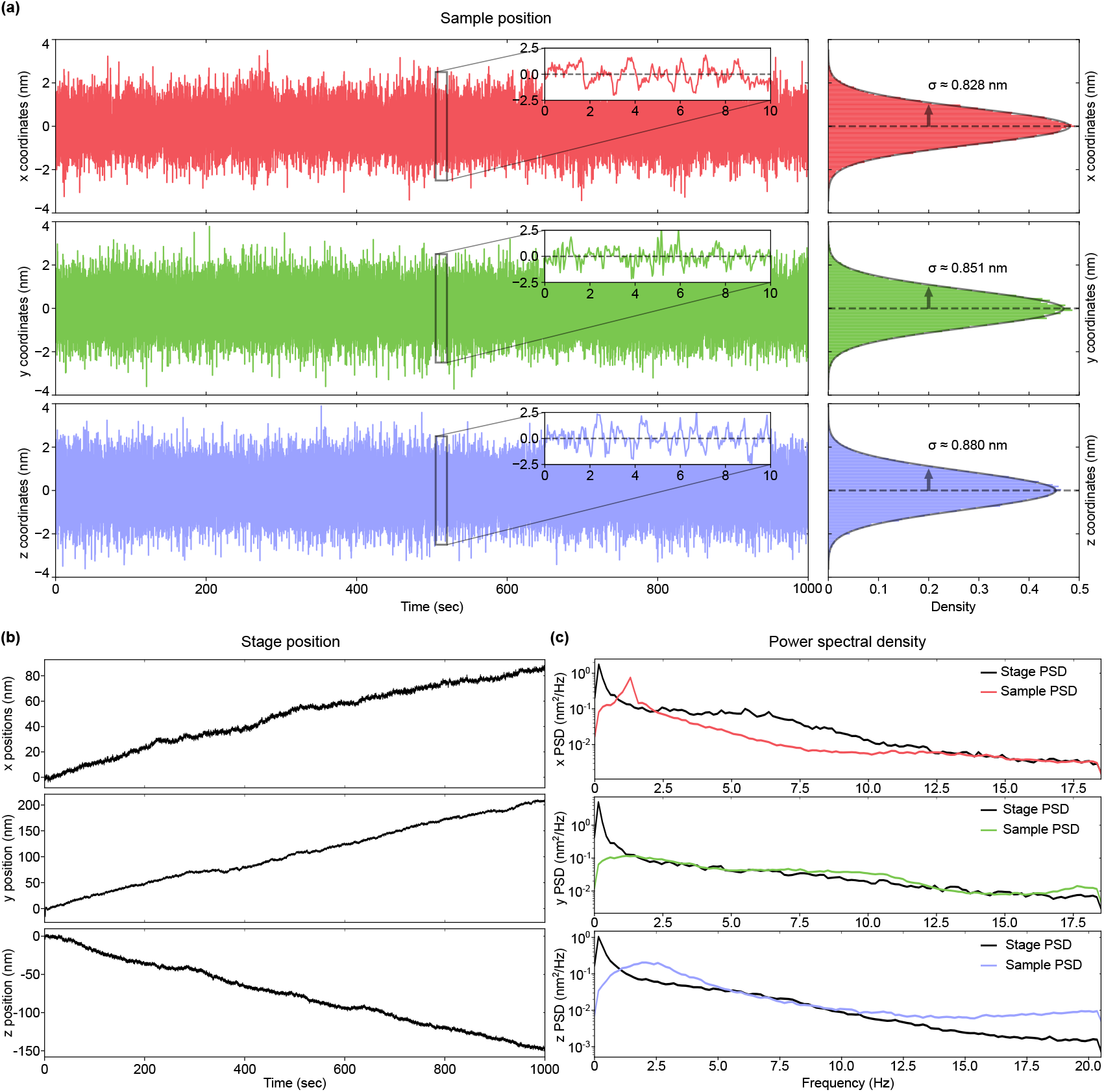
Stabilization results. (a) Sample position as monitored by the stabilization system and the corresponding histograms showing the distribution of the sample position across time. The parameter σ is the standard deviation of the Gaussian fit to the histogram. (b) Corresponding position of the piezoelectric stage for each axis. (c) Power spectral density (PSD) of the stabilized sample position (coloured) and the stage position (black).

As a control measurement, we performed long-term confocal imaging of the gold particle sample with the stabilization system running. Specifically, we removed the bandpass filter from our custom confocal and image scanning microscope to record reflection images in descanned mode using a single-photon avalanche diode (SPAD) array detector (35). Subsequently, we conducted time-series experiments utilizing our microscope control software, BrightEyes-MCS (36). We calculated the lateral drift in the confocal images by phase correlating each image with a reference image taken at the start of the measurement. With the stabilization software running, we observed a drift of (−0.8 ± 1.2) nm and (−0.6 ± 0.9) nm (mean ± standard deviation) in the x and y direction, respectively, over the course of 4 hours. Although these values exceed the sub-nm precision calculated by the stabilization system, these results may be expected given the additional uncertainty introduced by the galvanometric scan system or other drift components in sections of the beam path that are not common with the stabilization path. With the stabilization system turned off, we measured a lateral drift of more than 500 nm on the x-axis and more than 900 nm on the y-axis, with additional drift in the axial direction, moving the sample out of focus (Fig. 4 (b) & (d)).

**Fig 4.**
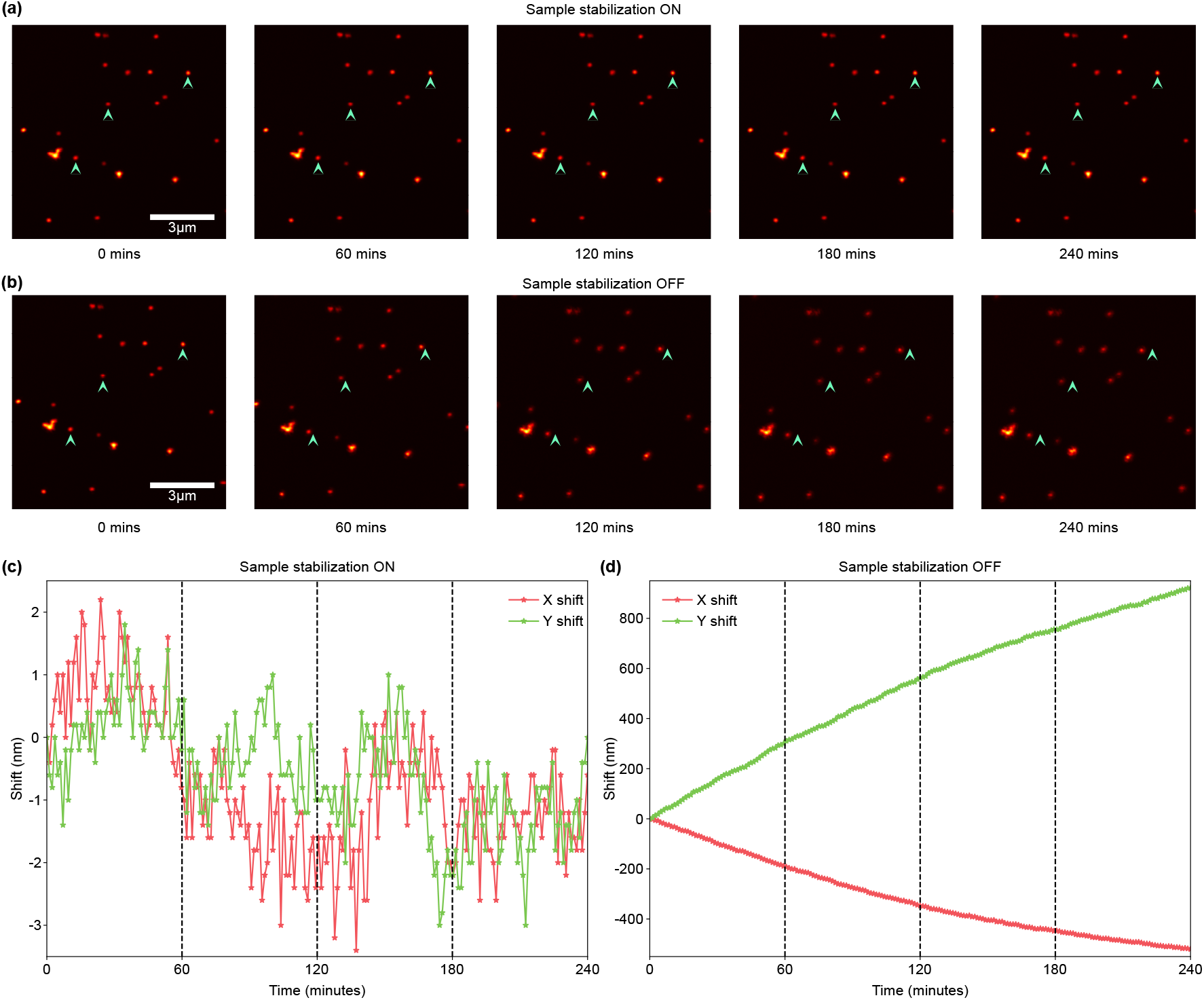
Confocal images acquired with and without active stabilization. (a) Subset of a time series taken with the active stabilization on. The sample consisted of gold nanoparticles attached to the coverslip surface. Images were acquired every minute for four hours. With the stabilization system on, there is no visible drift. (b) The same measurement with the stabilization system off shows a clear drift in both the lateral and axial directions over the course of four hours. (c, d) Measured drift in each image with respect to the start of the time-lapse, calculated by phase-correlation, with the stabilization system on (c) and off (d)

For both lateral and axial drift calculations, we prioritized lightweight algorithms over slower, more precise alternatives (34). Consequently, the update speed of our system is not limited by the software but by the rate at which the piezo stage controller can process movement commands, here 20 Hz for each axis. For applications where speed and accuracy are less critical, such as time-lapse measurements in confocal or widefield microscopy, alternative algorithms may be preferable. For instance, the local gradients method is well-suited to sparse particles whose images are radi-ally symmetric. However, for denser particle distributions or larger, non-radially symmetric structures, other methods, such as phase correlation, will perform better. Similarly, when only axial stabilization is required, analyzing the reflection of a beam off a cover slip can be advantageous, as it eliminates the need for fiducial markers in the sample. Although we have not implemented these alternatives here, the open-source code can be easily adapted to accommodate these custom scenarios.

In summary, we present an active 3*D* stabilization system for microscopy imaging, utilizing real-time monitoring of infrared back-scattering from fiducial markers within the sample. We capture and analyze widefield images at a rate of about 60 Hz and update the stage position at a rate of 20 Hz. Our approach yields sub-nanometer precision across all three spatial dimensions for several hours. The optical system is compact and portable, facilitating easy installation and transfer between different microscopes. The software is open-source and runs entirely on the CPU, removing the need for specialized hardware, such as a high-end GPU-equipped computer. We anticipate that these advantages will make super-resolution microscopy, as well as time-lapse confocal and widefield microscopy experiments, more accessible.

## Code availability

The Python code has been deposited on GitHub: https://github.com/VicidominiLab/Active_stabilization.

## Acknowledgements

This research was supported by: the European Research Council, BrightEyes No. 818699 (S.P., G.V., E.S.); the European Union’s Horizon 2020 research and innovation program under the Marie Sklodowska-Curie Grant Agreement No. 890923 (SMSPAD) (G.V., E.S.); PNRR MUR project PE00000023 – NQSTI and the European Union’s Horizon 2020 Research and Innovation Programme under Grant Agreement No. 881603 Graphene Flagship (G.V.).

## Supplementary Information

**Table S1.**
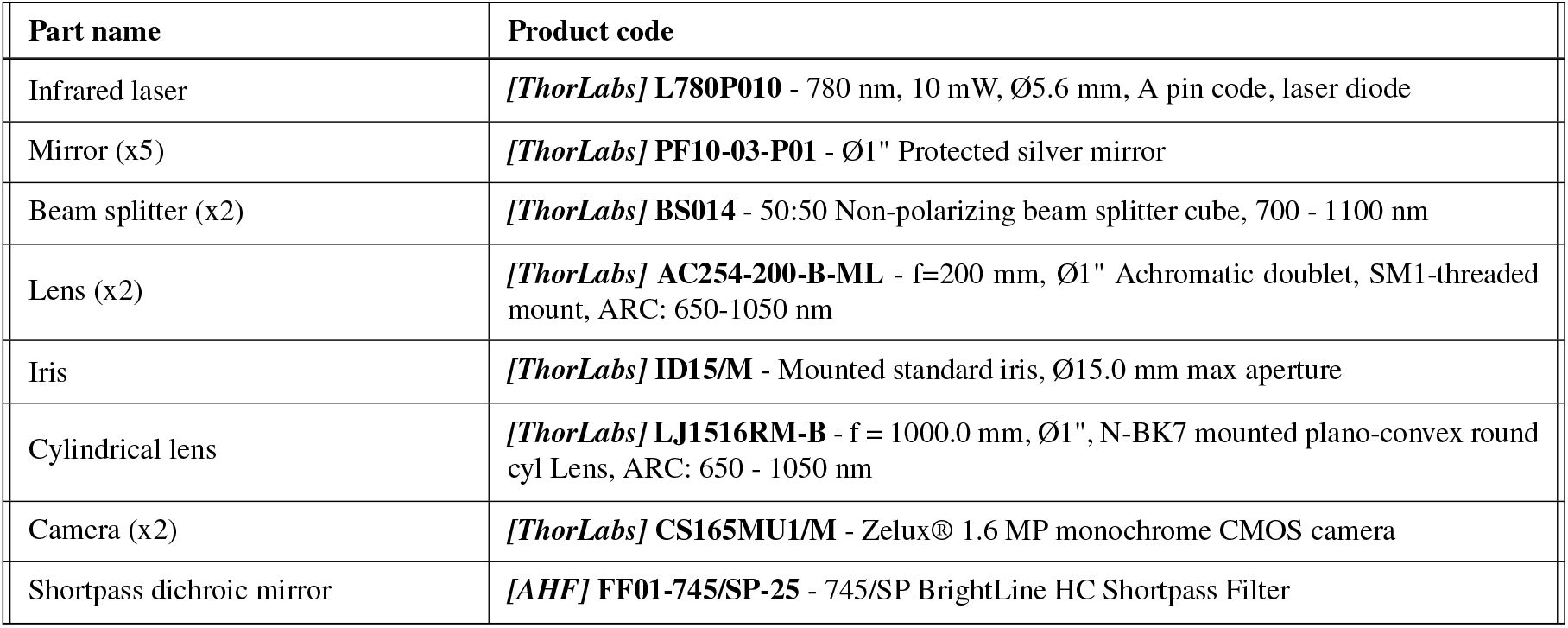
List of essential components required for building the stabilization system.

**Table S2.** List of additional (optional) components that may be used for building the stabilization system.

| Part name | Product code |
| --- | --- |
| Posts | *[ThorLabs]* **RS2P/M** - Ø25.0 mm Pedestal Pillar Post, M6 Taps, L = 50 mm |
| Mirror Mounts | *[ThorLabs]* **KM100** - Kinematic Mirror Mount for Ø1" Optics |
| Cage Plates | *[ThorLabs]* **CP35/M** - 30 mm Cage Plate with Ø1" Double Bore |
| Cage Rods | *[ThorLabs]* **ER3-P4** - Cage Assembly Rod (Variable size) |

